# The impact of age and education on phonemic and semantic verbal fluency: Behavioral and fMRI correlates

**DOI:** 10.1101/2021.01.14.426642

**Authors:** Rochele Paz Fonseca, Karine Marcotte, Lilian C. Hubner, Nicolle Zimmermann, Tânia Maria Netto, Bernardo Bizzo, Thomas Döring, J. Landeira-Fernandez, Emerson L. Gasparetto, Yves Joanette, Ana Inés Ansaldo

## Abstract

The purpose of this study was to examine the impact of age and education on the neural and behavioral correlates of verbal fluency. Forty-eight healthy adult participants were included: high-educated young and elderly, low-educated young and elderly. Participants performed semantic and phonemic and a control task during fMRI scanning. The phonemic fluency data showed an education effect across age groups. As for the semantic fluency data, there was an education effect only in young participants. The second-level fMRI results showed, in phonemic fluency, a main effect of age in the left posterior cingulate, superior temporal gyrus (STG) and right caudate, whereas the main effect of education involved activation in the right semantic fluency, there were a main effect of age in the left paracentral lobule and posterior cingulate, a main effect of education in the left claustrum and an interaction in the right claustrum and STG and the hippocampus bilaterally.

## 1. Introduction

Verbal fluency (VF) tasks have traditionally been administered as a clinical neuropsychological paradigm to assess linguistic, mnestic and/or executive abilities; they are also among the most widely used linguistic fMRI paradigms (1–4). The cognitive components assessed by this paradigm include executive abilities, such as initiation, inhibition, planning, updating and shifting; verbal long-term memory (word knowledge), both semantic and phonological; and lexical-semantic linguistic processes (5,6).

Since the end of the decade, neuroimaging studies have also been conducted using VF with healthy adults to characterize its neural correlates in an intact brain. Nevertheless, most such studies investigated brain activation during VF tasks in younger high-educated adults (4,7). Fewer studies have analyzed the neural correlates of VF in healthy elderly adults (5,8–11). Among the insufficiently analyzed results reported on the effect of aging on brain dynamics during VF, researchers found different patterns of recruitment of the left frontal lobe and more widespread activation in elderly people (11), which corroborates HAROLD (Hemispheric Asymmetry Reduction in Older Adults) model (12) which points to a dedifferentiation process occurring between the hemispheres, with increased participation of the right hemisphere compared to young adulthood and CRUNCH (Compensation-related utilization of neural circuits hypothesis) model (13) that argues that the engagement of more neural circuits than in younger adults could be explained by declining neural efficiency.

Comparing young and elderly participants’ performance in a VF task, Marsolais et al. (9) found a decrease in the functional integration of speech production networks and a significant interaction between task demand (comparing performance with more and less demanding semantic and orthographic stimuli) and age regarding functional connectivity in the anterior and posterior subnetworks of the VF network. Thus, age and education seem to play an important role both in behavioral performance and in the neural activity associated with language and other cognitive tasks, including VF; task demands also determine the patterns of brain activation.

Another issue with past studies using VF fMRI tasks is that almost all of them assessed participants with high levels of education. The brain modifications associated with educational experience have hardly been studied. Dehaene et al. (14) reviewed studies of illiterate adults and concluded that literacy reorganizes the brain to accommodate a novel cultural competence; the ventral occipito-temporal pathway and the right hemisphere are among the brain areas affected. Education can also be seen as one of the main components of cognitive reserve in aging (15). Evidence shows that brain damage may be overridden by the impact of education, as illustrated by the finding that different VF clusters in brain-damaged patients were better explained by level of education rather than by the lesion itself (16). Education was initially explored as a covariant in neuroimaging studies of aging, such as Nagels et al. (17). Nevertheless, the lower educational levels in most of these studies can still be considered as relatively high for some cultures and countries, apart from the wider range of educational levels in the sample recruited by Kim et al. (18) for a PET paradigm, where the low-educated people had less than six years of schooling.

Regarding methodological issues, fMRI studies have mainly used VF tasks that differ considerably from the procedures used in a standard clinical neuropsychological administration. First, although overt VF tasks have become more common (for a review, see Wagner et al. (1), more studies have been based on covert or silent paradigms (18–20). The main disadvantage of this procedure is that it includes errors among accurate answers (7). Additionally, even when the paradigm involves overt naming, one of the most frequent choices is an experiment-paced method (e.g., Abrahams et al. (21), in which the examinee recovers each word according to a fixed rhythm and not based on his/her self-monitoring. Another methodological issue relates to the duration of VF fMRI tasks, which are usually short, about 10 to 20 seconds (Allen and Fong, 2008; Birn et al., 2010; Wagner et al., 2014); only a few studies have used tasks lasting 60 seconds or more (e.g., Abrahams et al., 2003; Marsolais et al., 2014). Moreover, past studies were generally conducted with a block-design paradigm, which makes it impossible for each word to be treated as an event (7); the study by Marsolais et al. (9) is an exception in this respect. Regarding the fMRI system, of 26 fMRI studies with VF tasks, only 8 were conducted in a 3T scanner (1). There are also concerns related to the control tasks, which range from resting conditions to cognitively dissimilar control tasks, such as repetition of the same word, naming syllables or forward counting (1,20). The use of a month-listing task has been validated and is useful as a control automatic task (7). In general, there are not sufficient studies investigating both phonemic and semantic paradigms, both of which are very valuable tools in the clinical neuropsychology routine.

Surprisingly, despite the widespread interest in the effects of age and education on human cognition in behavioral studies of VF abilities, which generally report a negative impact of low education and decreased expressiveness as a result of aging (22–25), as far as we know no functional neuroimaging studies have yet investigated the possible impacts of age and education on semantic and phonemic VF and their neural correlates by comparing younger to older and better-educated to less-educated participants.

Therefore, working toward a better understanding of the effects of aging and educational background on brain organization and cognition, the purpose of this study is twofold: (1) to verify if age, education, or an interaction between these factors affects phonemic and semantic fMRI VF; and (2) to examine the neural correlates in brain activation or deactivation, if any, of age, education or their interaction. To do this, we used a recently developed mixed-design fMRI VF task (9). We also sought to analyze the relationship between brain activation or deactivation and VF performance. The innovative features of Marsolais et al.’s paradigm include the following: longer and thus more representative of clinical gold standards (90 seconds); overt; self-paced; alternate semantic and phonemic cues; control task involving automatic VF in naming months; and words considered as events. Furthermore, for the first time, we applied this hitherto little-used paradigm, on which only two studies had been based (9,26), to analyze the impacts of both age and education on underlying cognitive abilities.

Some predictions were made. First, we expected that education would be the most prominent variable affecting VF performance in both phonemic and semantic tasks. In association with high-functioning aging, more diffuse and right-lateralized brain activation should be found in elderly participants. More intense deactivation of language- and executive-related regions should be found for low-educated adults, insofar as evoking words would be more difficult for them, while greater activation should be found for higher-educated participants. Finally, we expected to see a continuum of activation and deactivation, both fostering better performance in VF tasks, related to the level of complexity and executive function recruitment, as follows: high-educated young adults (HY) would show more deactivation due to their greater facility in performing the tasks, followed by high-educated elderly adults (HE) and low-educated young adults (LY), who would present a mid-range activation/deactivation pattern, while in low-educated elderly adults (LE) some executive control regions would be activated more further to their attempts to perform better.

## 2. Methods

### 2.1 Participants

Sixty-seven participants were first recruited for this study based on their age and education. The final sample contained 48 participants, who were divided into four groups (n = 12 each) according to their age and education level: (1) LY, (2) HY, (3) LE and (4) HE. Low- and high-educated groups were matched by age, and older and younger groups were matched by education, socioeconomic level, and psychological test scores (intelligence, dementia and depression). All four groups were matched by gender; there were 6 men and 6 women in each group. Younger participants ranged from 19 to 40 years old, while elderly participants were aged 60 to 77 years old. Low-educated adults had received from 5 to 10 years of formal education, whereas high-educated participants had 12 years or more of formal education. Detailed demographic characteristics and neuropsychological testing scores of the four groups are reported in Table 1.

**Table 1.**
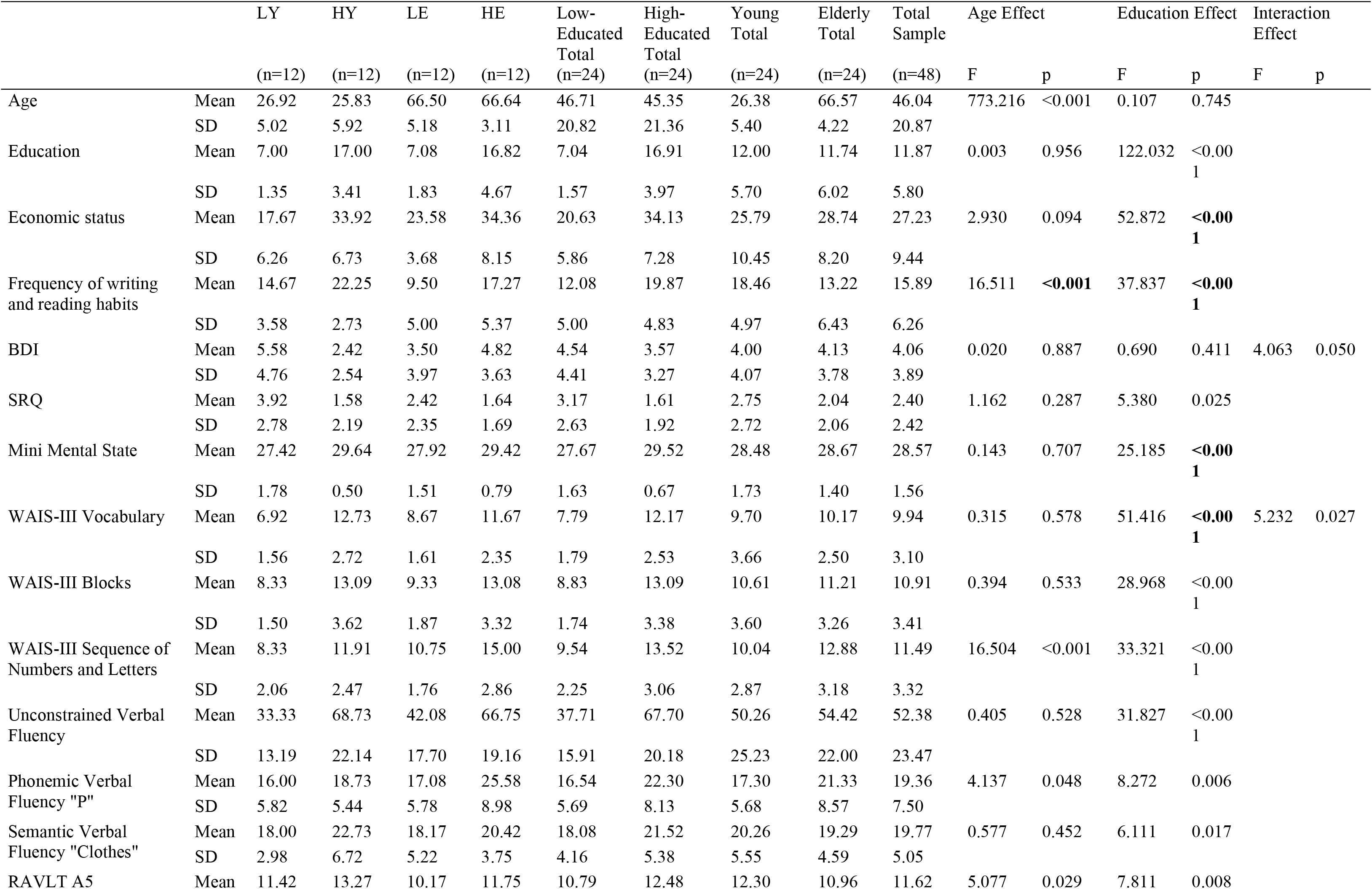
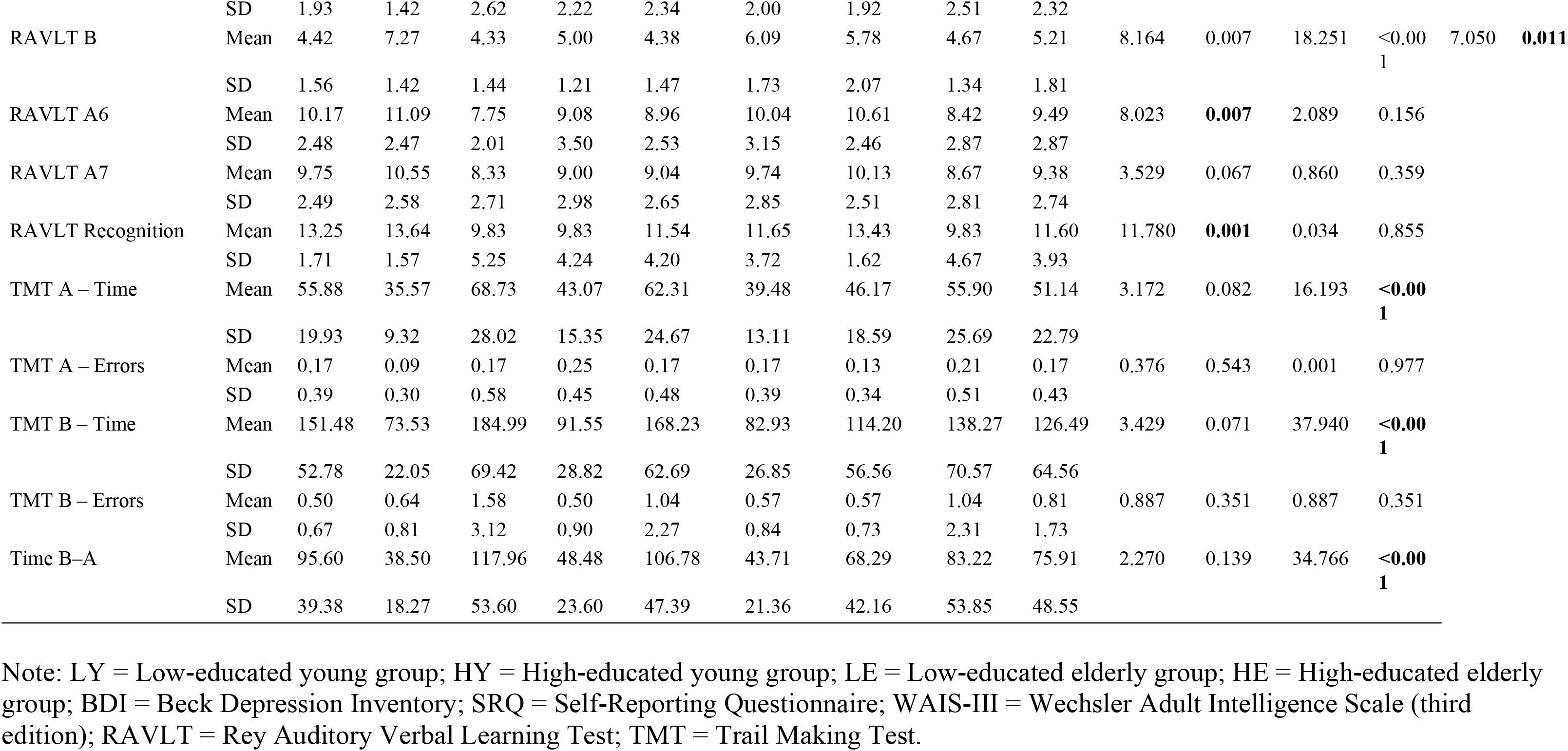
Sociodemographic characteristics. neuropsychological screening scores and behavioral results per group

Inclusion criteria were (1) participants should be monolingual native speakers of Brazilian Portuguese; and (2) right-handed prior to the study, according to a cut-off score of 80% right dominance in the Brazilian Portuguese version of the Edinburgh Inventory test (27). Exclusion criteria were (1) self-report of neurological or psychiatric illness; (2) use of medications with psychotropic effects; (3) abnormal or uncorrected visual or auditory acuity; (4) identification of brain tissue abnormality on structural MRI; and (5) presence of Mild Cognitive Impairment (MCI) or dementia prior to the study. To screen for signs of MCI or dementia, the Mini Mental State Examination (27–30) was administered to all participants; for low-educated individuals, a minimum score of 24 was required, whereas a minimum score of 28 was required for high-educated participants. To screen for depression signs, the Beck Depression Inventory (BDI) (31,32) was administered, and participants with a score higher than 9 were excluded. Finally, the Vocabulary and Block design dyad of the Wechsler Adult Intelligence Scale (WAIS-III) (33,34) was administered to exclude any participants with intellectual impairments.

A total of 19 participants were excluded: 3 due to technical difficulties with audio recordings or missing imaging data, 9 because of MRI results compatible with neurovascular diseases (images compatible with stroke, traumatic brain injury or brain tumor) or claustrophobia, and 7 because they misunderstood or did not complete the task, due to lower educational level. All participants were recruited on a voluntary basis in the community, at the university or at work. This research project was approved by the Ethical Committee of the Pontifical Catholic University of Rio Grande do Sul. Written informed consent was obtained from all individuals before they took part in the experiment.

### 2.2 Procedure, design and paradigm

All participants were individually evaluated at the Clínica de Diagnóstico por Imagem, Rio de Janeiro, Brazil, by three trained neuropsychologists. A neuroradiologist first performed an anatomical screening for all participants to check for exclusion criteria. When selected for the study, they performed three training VF tasks, based on the tasks in the *Protocole Montréal d’Évaluation de la Communication* (2), adapted for use in Brazil (35). First, participants were asked to say as many words as possible for two and a half minutes, without any constraint other than not using proper nouns or numbers (free VF). Then, they were asked to name as many words as possible belonging to a semantic category (fruits) for two minutes (semantic fluency). Finally, they had to name as many words as possible beginning with the letter F for two minutes (phonemic fluency). They were selected for the fMRI procedures only if, as well as meeting all standard prerequisites for an fMRI study, they were able to complete all three training tasks. Then each participant was scanned with MRI and fMRI sequences, while he/she performed the experimental tasks.

#### 2.2.1 Design and paradigm

Participants performed an overt, self-paced VF task. A mixed design was used. Specifically, data were collected in blocks of different VF tasks, but the words were analyzed as events, thus using a mixed design. The whole experiment was conducted in eight epochs, in which we alternated a resting phase (10 seconds), the control task (90 seconds), a resting phase (10 seconds), and the semantic or phonemic fluency task (90 seconds). Thus, each epoch lasted 200 seconds and was part of a unique run lasting 1,600 seconds. Each phase was presented visually in black and white at the center of the screen while the task lasted. The eight epochs started with a “?”centered on the screen (resting phase), followed by “months” (control task), “?” again (resting phase), and, finally, “letter” or “category” (phonemic or semantic fluency task). The experimental tasks were counterbalanced across subjects, in four different orders. The design of an epoch is represented in Figure 1.

**Figure 1.**
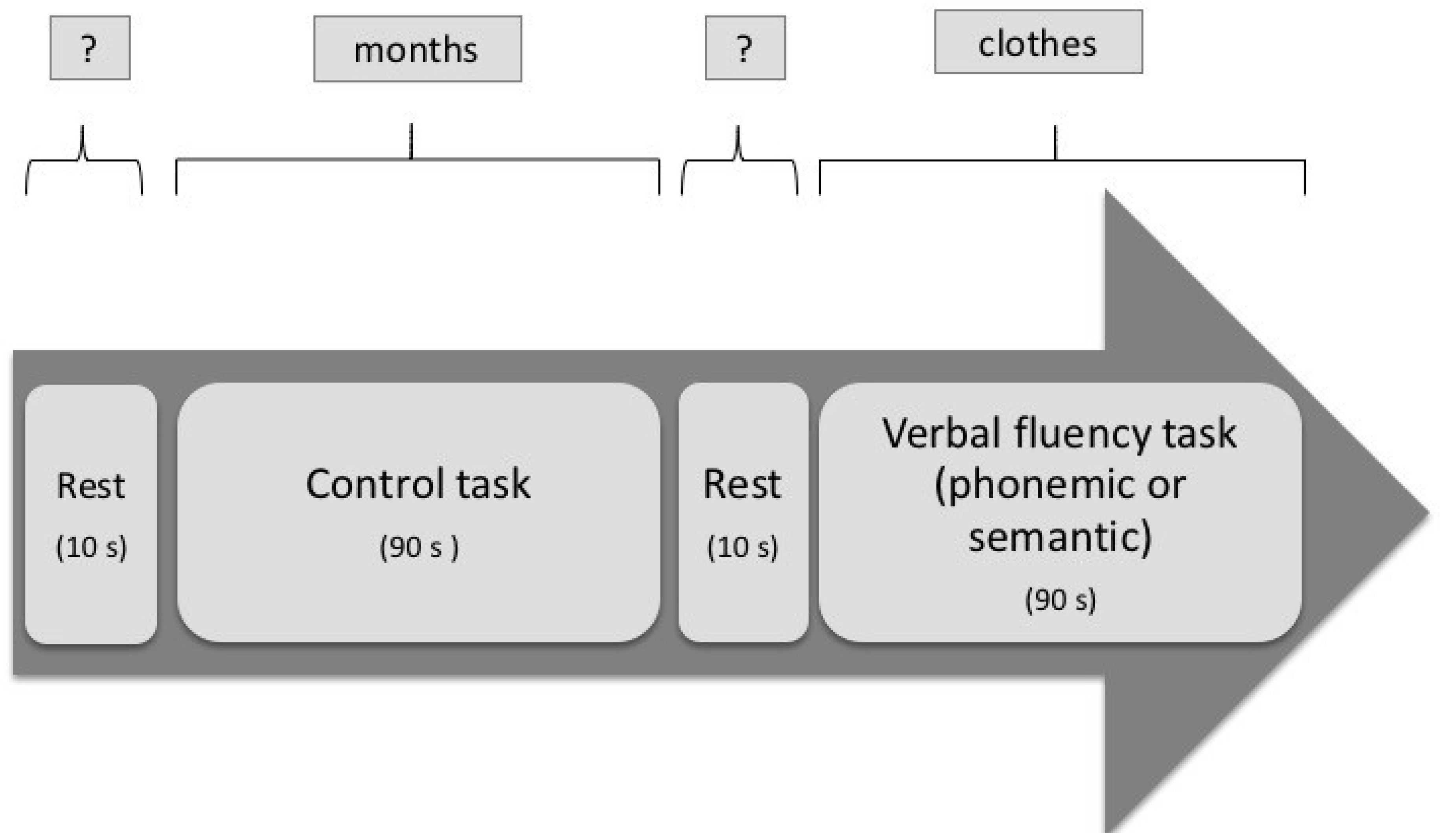
Illustration of an epoch of the mixed-design verbal fluency paradigm for fMRI

A training session preceded imaging acquisition. The training task consisted of a task similar to the experimental one, but with different stimuli: the letter F in the phonemic condition and “fruits” in the semantic condition. Participants were instructed to generate as many words as possible belonging each criterion. The experimental fMRI VF task consisted in naming the semantic categories of animals, clothes, sports and vegetables, while the phonemic task required participants to name words starting with the letters P, M, L and V. These experimental tasks had previously been pilot-tested outside the scanner in a separate group of 20 healthy adults, with successful results. The control task consisted of naming the months of the year consecutively, beginning with January, continuing to December and restarting with January.

### 2.3 fMRI acquisition

Brain images were acquired with a 3T scanner (Verio, Siemens Medical, Erlangen Germany) using a 12-channel head matrix coil. Before the fMRI scans, structural images were obtained in a sagittal plane using a T1-weighted 3D MPRAGE sequence (TR/TE = 2530/3.43 ms, FOV = 256 mm, matrix size 192 × 256 × 128, voxel size 1.3 × 1.0 × 1.3 mm^3^, slice thickness = 1.3 mm; acquisition = 128 slices in the axial plane, with a distance factor of 50%, so as to scan the whole brain, including the cerebellum). Functional data were obtained using a gradient echo EPI sequence in the transversal plane (TR/TE = 2000/30 ms, FOV = 215 mm, matrix size 122 × 122 × 28, voxel size = 3.4 × 3.4 × 5.0 mm, interleaved acquisition = 28 slices with a distance factor of 0%). The transversal plane was aligned along the AC-PC line. Finally, 800 volumes were acquired (TR = 2; TE = 30 ms). In order to reduce head movements, the subject’s head was restrained by foam pads inside the head coil.

Visual stimuli were presented via E-prime (Psychology Software Tools) from a computer onto a screen at the head of the bore, and were visible in a mirror attached to the head coil. Participants’ verbal responses were recorded using an MRI compatible optical dual channel active noise-canceling microphone system (FoMRI II, Optoacoustics, Or-Yehuda, Israel), also mounted on the head coil. The microphone was connected to an optical unit in order to record the responses with a coupled workstation. In this way, almost all scanner noise could be eliminated to achieve greater intelligibility when transcribing and analyzing verbal responses.

### 2.4 Data analyses

#### 2.4.1 Behavioral analyses

All evoked words were transcribed and analyzed as correct or incorrect answers. Correct answers included words that responded to the criterion and were non-perseverative, including synonyms. A total semantic fluency score was generated by adding the four scores for each category, and the same was done to generate a phonemic VF score. In order to avoid a large difference between the quantity of events of both semantic and phonemic VF and of the control task, and thus allow a comparison between behavioral and neuroimaging data, only the middle 10 seconds of the months fluency task in each epoch was summed to generate a total months task score for the eight epochs. These three VF accuracy scores were entered in a two-way ANOVA (age and education factors), in SPSS 18 for Windows, with p < 0.05.

#### 2.4.2 fMRI analyses

##### 2.4.2.1 Whole-brain analyses

Raw data visualization and preprocessing were conducted with Matlab 7.11 (MathWorks, Natick, MA) using standard procedures in SPM8 software (Wellcome Department of Imaging Neuroscience, UK; http://www.fil.ion.ucl.ac.uk/spm). Preprocessing involved slice timing, realignment (to account for head movements), coregistration, segmentation, normalization and smoothing. The segmented images were corrected and spatially normalized to the Montreal Neurological Institute’s standard. The normalized functional images were spatially smoothed with an 8-mm Gaussian filter.

For further analysis, each correct word’s onset was precisely established in milliseconds using SoundForge 10 (Sony Pictures Digital, Inc.). First-level analyses were done using the general linear model in SPM8. For subject-level analysis, statistical parametric maps were obtained for each participant, by means of linear contrasts applied to the parameter estimates for the studied events. Therefore, for each individual, t-contrasts of the experimental conditions versus control condition were calculated for each voxel (semantic VF – months; phonemic VF – months). Separate t-tests against the null hypothesis that there was no change in activation were performed for each contrast and each group. This resulted in contrast maps used for group-level analysis. To quantify left or right lateralization for each group, a lateralization index was calculated for each contrast. The total of activated pixels in the left and right hemispheres was obtained, calculating an asymmetry index with the formula (L–R)⁄(L+R), considering activation to be left hemisphere (L) dominant if this score is higher than 0.20 and right-dominant (R) if the index is lower than –0.20 (36). Using this formula, a positive value indicates higher activation in the left hemisphere whereas a negative value indicates a lateralization in the right hemisphere.

Second-level group analyses were conducted in SPM by means of t-tests and ANOVAs. By employing a one-sample t-test, averages were calculated for each condition and for the four groups. To compare the groups and explore a possible interaction between age and education, a full factorial analysis was done for each contrast examined in a single analysis (phonemic VF – months and semantic VF – months). Each factor had two levels: age (older and younger) and education level (low and high). When a main effect or an interaction was found, a two-sample t-test was conducted to analyze the following subtractions: older versus younger, younger versus older, low- versus high-educated, high-versus low-educated. As is commonly done in the literature, and since random effects analyses may be fairly conservative in fMRI data (37), a threshold of p < 0.001 uncorrected was used. Contrasts were performed with a cluster size of at least 10 voxels (k > 10).

Considering the final sample size, when comparing age and education groups, the only possible contrasts to be included as dependent variables were semantic – control task and phonemic – control task (it was not possible to include VF modality or different frequency levels of productivity as independent variables, since they were counterbalanced in each modality). Although our initial sample size was acceptable, the sample for each age and education group (n = 12) was so small that additional subgroup analysis would inflate the chances of false positive or negative errors. Moreover, we would be testing a hypothesis that had not been sufficiently explored in neuroimaging studies with larger samples, which is not a positive scenario for subgroup analyses (Sun et al., 2010).

##### 2.4.2.2 Region of interest analyses

Finally, in order to better explore the observed effects in between-subject comparisons found with the ANOVAs, a complementary analysis was conducted. Aiming to obtain region of interest (ROI) data from each participant’s images, the percentage of signal change was extracted for each region where a main effect of age or education or an interaction between the two factors was observed, using MarsBaR 0.42 (http://marsbar.scourceforge.net) software for ROI analyses. In this way, real activations could be identified for each group. To further investigate the relationship between activation or deactivation and VF performance, the percentage of signal change of each ROI for each participant was correlated with the individual performance scores for phonemic and semantic VF tasks through a Pearson linear analysis.

## 3. Results

### 3.1 Behavioral data

Sociodemographic characteristics, neuropsychological screening scores and behavioral results for each group are reported in Table 1. Subgroup analysis of interaction effects with post hoc analysis with the Bonferroni procedure showed that the four groups had similar scores on the BDI. As expected, LY individuals scored significantly lower on the WAIS-III Vocabulary subtest than HY (p < 0.001) and HE individuals (p < 0.001), but had similar scores to the LE group (p = 0.304). On the same subtest, the HY group scored significantly higher than the LE group (p < 0.001), but not the HE group (p = 0.798). HE individuals had significantly higher scores than LE participants (p = 0.008). Significant differences in performance were also encountered on the RAVLT B, on which the HY individuals outperformed the LY (p < 0.001), LE (p < 0.001) and HE participants (p = 0.001). Finally, on the semantic VF task, LY participants scored lower than HY (p < 0.001) and HE participants (p = 0.001), and HY individuals performed better than the LE group (p < 0.001). There were no age differences between the education groups (t(46) = –0.103, p = 0.918), and no education differences between the age groups (t(46) = 0.000, p = 1.000). We found an education effect on the economic status score (Higher Education > Lower Education), and main effects of age and education on the frequency of reading and writing habits. Descriptive data on the mean phonemic and semantic VF performance and on the control task (months) for each group are also reported in Table 1.

#### Post hoc Bonferroni correction (interactions)

After post hoc analysis with the Bonferroni procedure, the four groups presented similar scores on the BDI. As expected, LE individuals scored significantly lower on the WAIS-III Vocabulary subtest than HY (p < 0.001) and HE individuals (p < 0.001), but performed similarly to the LE group (p = 0.304). For the same subtest, the HY group scored significantly higher than the LE group (p < 0.001), but not the HE group (p = 0.798). In addition, HE individuals performed significantly better than the LE group (p = 0.008). Significant differences in performance were also observed on the RAVLT B score, such that HY individuals outperformed LE (p < 0.001), LE (p < 0.001) and HE individuals (p = 0.001).

### 3.2 fMRI data

After conducting a one-sample t-test in a within-group analysis, activated regions per group for the phonemic – months contrast (Table 2) and for the semantic –months contrast (Table 3) were found. During the phonemic VF task, all groups except the HE group activated the right cingulate gyrus among other regions. LY and LE had activated the left insula. For semantic fluency, both LY and HY groups activated the left caudate tail. In addition, HE participants activated a greater range of regions, mainly in the right hemisphere, such as the bilateral anterior lobe of the cerebellum, right posterior lobe of the cerebellum, and bilateral superior temporal gyrus (STG).

**Table 2.** Activated brain areas for Phonemic VF versus Months contrast per group, p < 0.001, uncorrected, k > 10

|  | x | Y | z | cluster | Z | x | y | z | cluster | Z |
| --- | --- | --- | --- | --- | --- | --- | --- | --- | --- | --- |
|  | Left Hemisphere |  |  |  |  | Right Hemisphere |  |  |  |  |
| Low-educated younger |  |  |  |  |  |  |  |  |  |  |
| Insula (13) | −28 | −2 | 26 | 97 | 3.57 |  |  |  |  |  |
| Limbic lobe, subgyral (24) | −22 | −16 | 40 | 97 | 3.54 |  |  |  |  |  |
| Cingulate gyrus (32/24) |  |  |  |  |  | 18 | 20 | 26 | 73 | 5.25 |
| Clastrum | −24 | 26 | 16 | 81 | 3.54 | 28 | 10 | 18 | 11 | 3.45 |
| High-educated younger |  |  |  |  |  |  |  |  |  |  |
| Cerebellum, posterior lobe |  |  |  |  |  | 28 | −54 | −48 | 35 | 4.04 |
| Postcentral gyrus (3) | −42 | −18 | 34 | 99 | 3.77 |  |  |  |  |  |
| Posterior cingulate gyrus (31) |  |  |  |  |  | 28 | −68 | 12 | 11 | 4.99 |
| Low-educated elderly |  |  |  |  |  |  |  |  |  |  |
| Insula (13) | −30 | 28 | 14 | 51 | 3.78 |  |  |  |  |  |
| Clastrum |  |  |  |  |  | 24 | 28 | 14 | 94 | 3.58 |
| Anterior cingulate gyrus (32) |  |  |  |  |  | 24 | 40 | 6 | 94 | 3.47 |
| High-educated elderly |  |  |  |  |  |  |  |  |  |  |
| Middle frontal gyrus (9) | −16 | 32 | 36 | 19 | 3.39 |  |  |  |  |  |

**Table 3.** Activated brain areas for Semantic VF versus Months contrast per group, p < 0.001, uncorrected, k > 10

|  | x | Y | z | cluster | Z | x | y | z | cluster | Z |
| --- | --- | --- | --- | --- | --- | --- | --- | --- | --- | --- |
|  | Left Hemisphere |  |  |  |  | Right Hemisphere |  |  |  |  |
| Low-educated younger |  |  |  |  |  |  |  |  |  |  |
| Caudate tail | −16 | −34 | 18 | 35 | 3.80 |  |  |  |  |  |
| High-educated younger |  |  |  |  |  |  |  |  |  |  |
| Caudate tail | −26 | −40 | 14 | 10 | 3.49 |  |  |  |  |  |
| Low-educated elderly |  |  |  |  |  |  |  |  |  |  |
| Cingulate gyrus (32) |  |  |  |  |  | 22 | 12 | 30 | 29 | 3.75 |
| High-educated elderly |  |  |  |  |  |  |  |  |  |  |
| Cerebellum, anterior lobe | −4 | −44 | −14 | 749 | 4.05 | 32 | −46 | −34 | 245 | 4.22 |
| Occipital lobe, lingual gyrus (19) | −10 | −64 | −6 | 749 | 3.83 |  |  |  |  |  |
| Middle frontal gyrus (9) | −4 | −50 | −28 | 749 | 3.71 |  |  |  |  |  |
| Parahippocampal gyrus (19) |  |  |  |  |  | 44 | −38 | −10 | 32 | 3.66 |
| Hippocampus |  |  |  |  |  | 38 | −28 | −14 | 32 | 3.26 |
| Cerebellum, posterior lobe |  |  |  |  |  | 20 | −84 | −28 | 16 | 5.20 |
|  |  |  |  |  |  | 38 | −66 | −40 | 47 | 3.38 |
| Cingulate gyrus (32) | −16 | 8 | 38 | 118 | 3.61 |  |  |  |  |  |
| Superior temporal gyrus (41) | −52 | −18 | −4 | 11 | 3.23 | 60 | −16 | 2 | 44 | 3.56 |

In summary, regarding the main regions involved, VF was associated with activation in the STG, cingulate gyrus, cerebellum, claustrum, insula and caudate tail. For phonemic fluency, the lateralization index was 0.53 for the LY group, 0.36 for the HY group, –0.57 for the LE group, and 1 for the HE group. Thus, in this letter-based task, all groups revealed left-hemisphere dominance, except for the LE group, which activated more right-hemisphere regions (see Figure 2). For semantic fluency, the lateralization index was 1 for both LY and HY groups, –1 for the LE group, and 0.68 for the HE group. Again, the LE group was the only one showing right-hemisphere dominance (see Figure 3).

**Figure 2.**
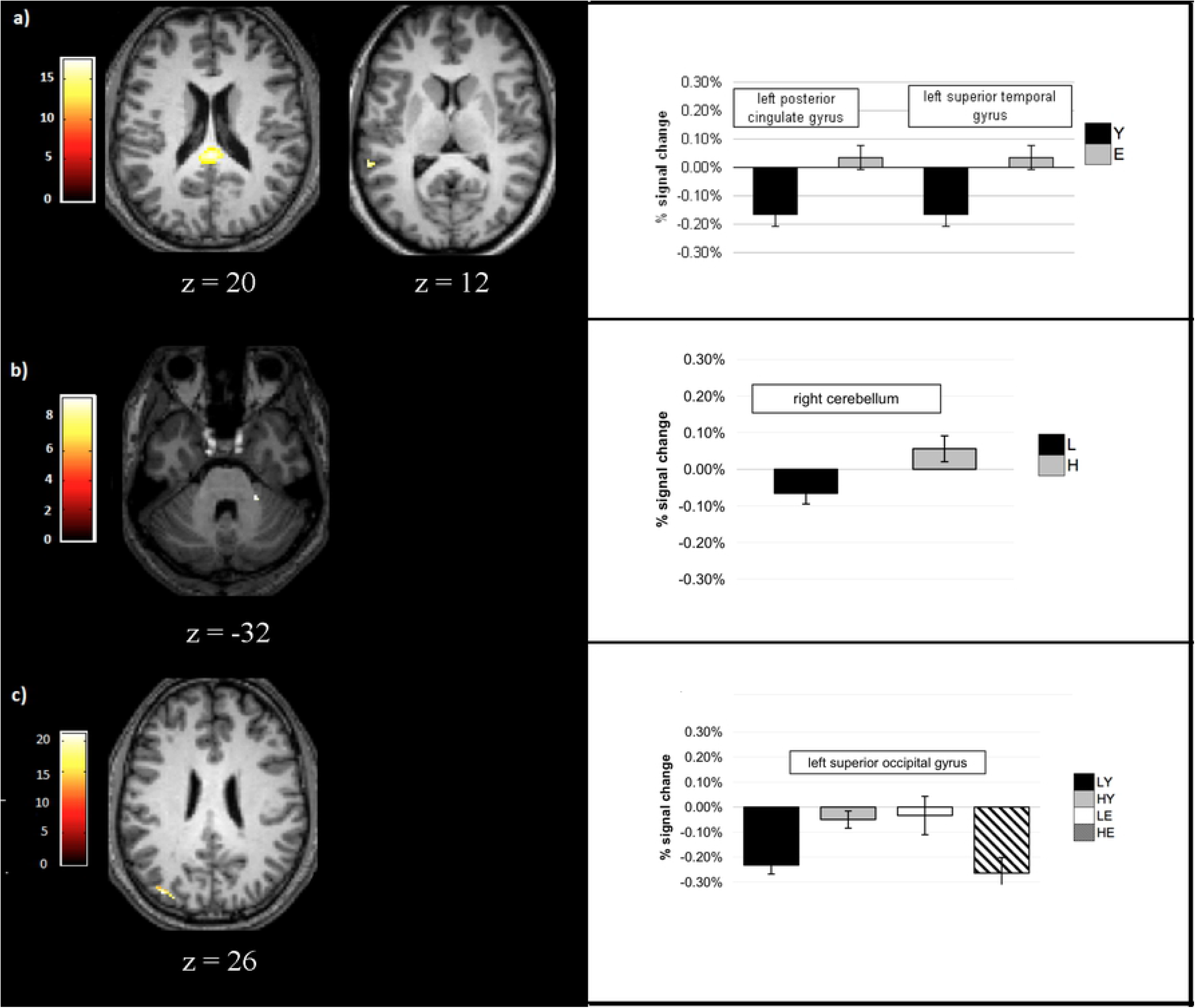
Patterns of activation changes in the phonemic fluency task for the (a) age effect, (b) education effect and (c) interaction between age and education (first column) and histograms for percentage of signal change per group in ROIs related to each effect Note: The activation in the cerebellum for the education effect (b) was observed with a less conservative threshold.

**Figure 3.**
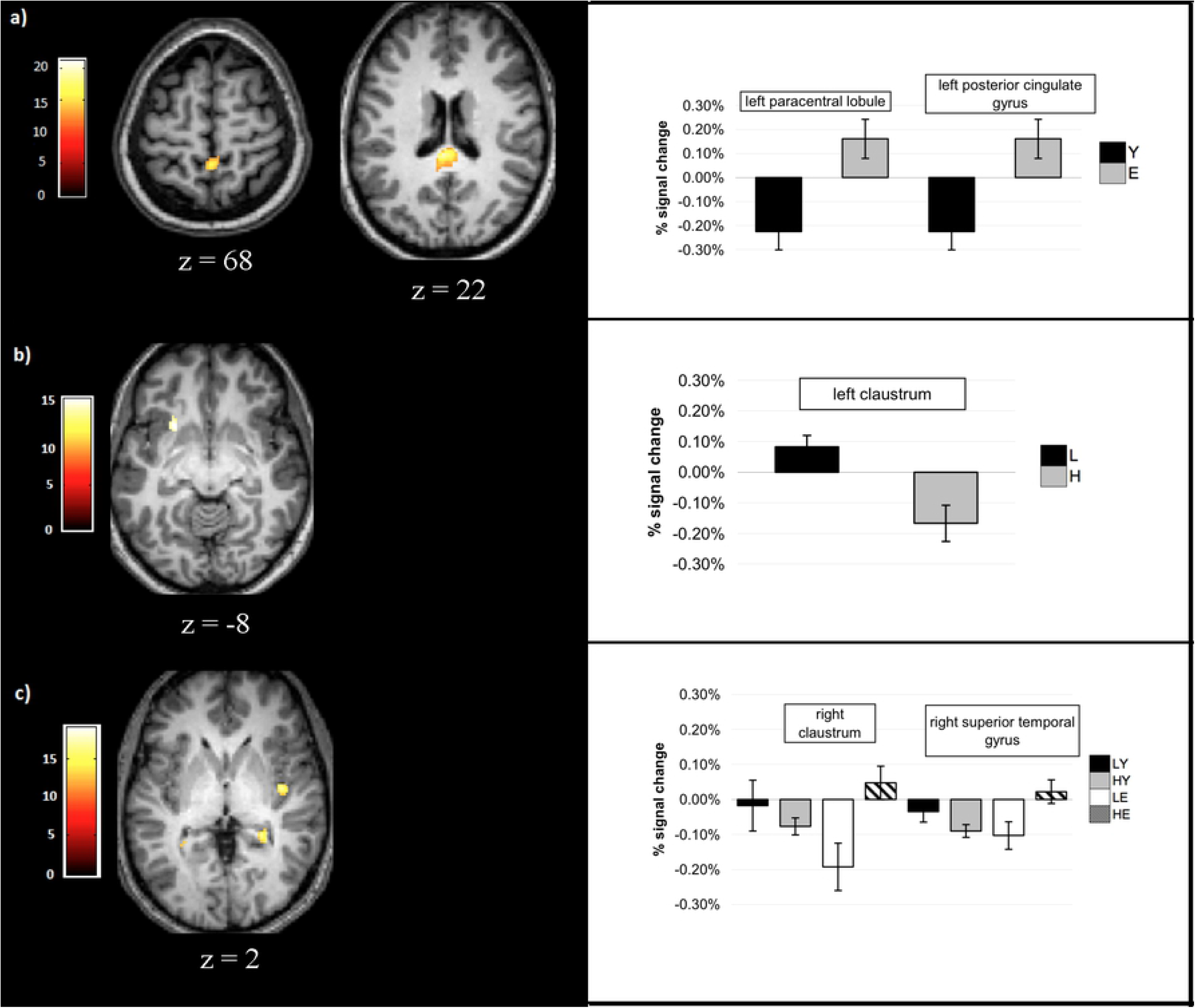
Patterns of activation changes in the semantic fluency task for the (a) age effect, (b) education effect and (c) interaction between age and education (first column) and histograms for percentage of signal change per group in ROIs related to each effect

fMRI second-level results showed a main effect of age for phonemic VF, manifested in the left posterior cingulate gyrus (PCG), left STG and right caudate, whereas a main effect of education was found in the right cerebellum. There was an interaction in the left superior occipital gyrus (SOG). For semantic VF, there was a main effect of age in the left paracentral lobule and PCG, whereas a main effect of education was observed in the left claustrum. An interaction between age and education was found in the right claustrum and STG, as well as in the hippocampus bilaterally (see Table 4).

**Table 4.**
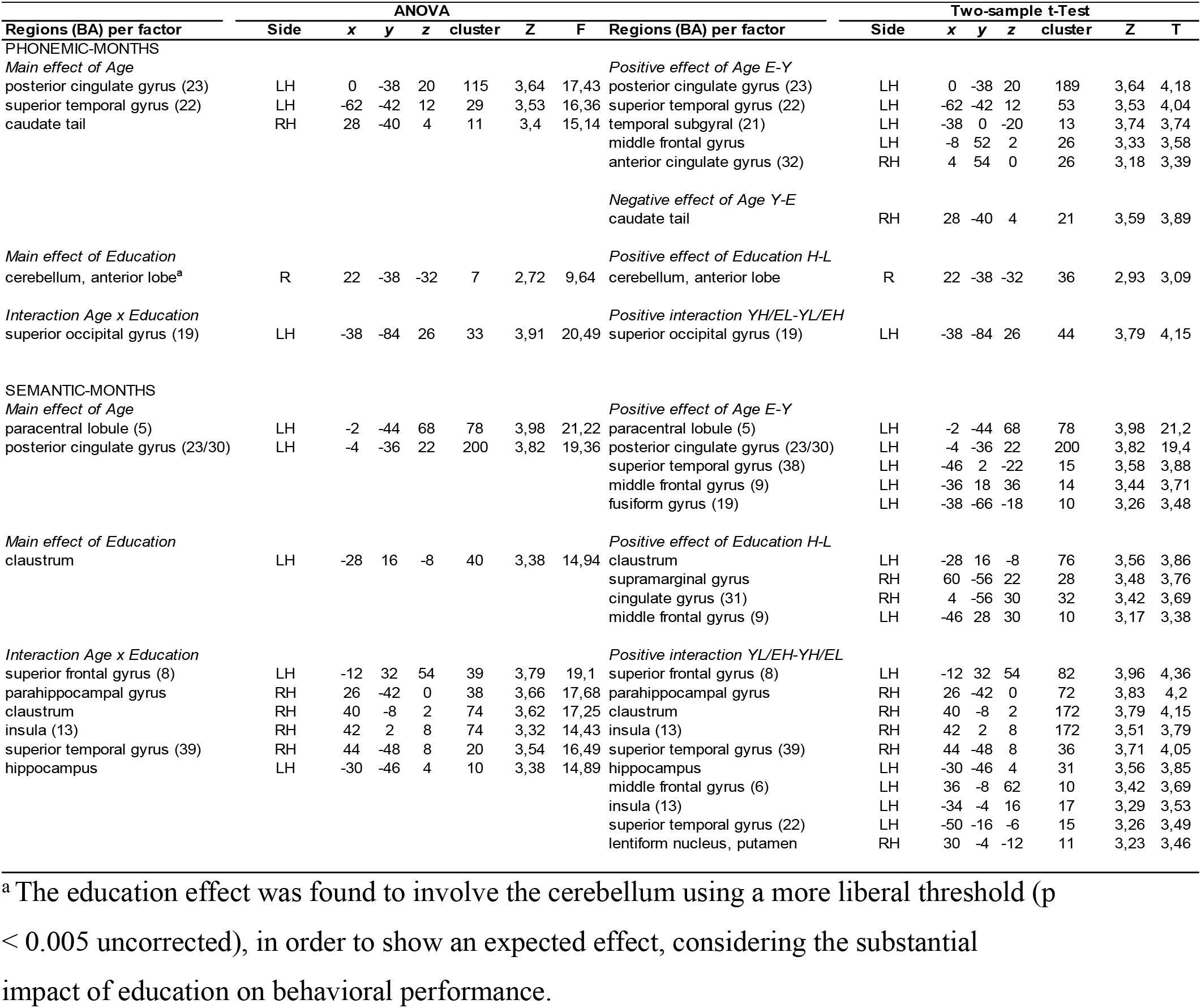
fMRI data from second-level analyses (full factorial analyses and two-sample t Tests for interpretation)

| ANOVA |  |  |  |  |  |  |  | Two-sample t-Test |  |  |  |  |  |  |  |
| --- | --- | --- | --- | --- | --- | --- | --- | --- | --- | --- | --- | --- | --- | --- | --- |
| Regions (BA) per factor | Side | x | y | z | cluster | Z | F | Regions (BA) per factor | Side | x | y | z | cluster | Z | T |
| PHONEMIC-MONTHS |  |  |  |  |  |  |  |  |  |  |  |  |  |  |  |
| Main effect of Age |  |  |  |  |  |  |  | Positive effect of Age E-Y |  |  |  |  |  |  |  |
| posterior cingulate gyrus (23) | LH | 0 | -38 | 20 | 115 | 3,64 | 17,43 | posterior cingulate gyrus (23) | LH | 0 | -38 | 20 | 189 | 3,64 | 4,18 |
| superior temporal gyrus (22) | LH | -62 | -42 | 12 | 29 | 3,53 | 16,36 | superior temporal gyrus (22) | LH | -62 | -42 | 12 | 53 | 3,53 | 4,04 |
| caudate tail | RH | 28 | -40 | 4 | 11 | 3,4 | 15,14 | temporal subgyral (21) | LH | -38 | 0 | -20 | 13 | 3,74 | 3,74 |
|  |  |  |  |  |  |  |  | middle frontal gyrus | LH | -8 | 52 | 2 | 26 | 3,33 | 3,58 |
|  |  |  |  |  |  |  |  | anterior cingulate gyrus (32) | RH | 4 | 54 | 0 | 26 | 3,18 | 3,39 |
|  |  |  |  |  |  |  |  | Negative effect of Age Y-E |  |  |  |  |  |  |  |
|  |  |  |  |  |  |  |  | caudate tail | RH | 28 | -40 | 4 | 21 | 3,59 | 3,89 |
| Main effect of Education |  |  |  |  |  |  |  | Positive effect of Education H-L |  |  |  |  |  |  |  |
| cerebellum, anterior lobe<sup>a</sup> | R | 22 | -38 | -32 | 7 | 2,72 | 9,64 | cerebellum, anterior lobe | R | 22 | -38 | -32 | 36 | 2,93 | 3,09 |
| Interaction Age x Education |  |  |  |  |  |  |  | Positive interaction YH/EL-YL/EH |  |  |  |  |  |  |  |
| superior occipital gyrus (19) | LH | -38 | -84 | 26 | 33 | 3,91 | 20,49 | superior occipital gyrus (19) | LH | -38 | -84 | 26 | 44 | 3,79 | 4,15 |
| SEMANTIC-MONTHS |  |  |  |  |  |  |  |  |  |  |  |  |  |  |  |
| Main effect of Age |  |  |  |  |  |  |  | Positive effect of Age E-Y |  |  |  |  |  |  |  |
| paracentral lobule (5) | LH | -2 | -44 | 68 | 78 | 3,98 | 21,22 | paracentral lobule (5) | LH | -2 | -44 | 68 | 78 | 3,98 | 21,2 |
| posterior cingulate gyrus (23/30) | LH | -4 | -36 | 22 | 200 | 3,82 | 19,36 | posterior cingulate gyrus (23/30) | LH | -4 | -36 | 22 | 200 | 3,82 | 19,4 |
|  |  |  |  |  |  |  |  | superior temporal gyrus (38) | LH | -46 | 2 | -22 | 15 | 3,58 | 3,88 |
|  |  |  |  |  |  |  |  | middle frontal gyrus (9) | LH | -36 | 18 | 36 | 14 | 3,44 | 3,71 |
|  |  |  |  |  |  |  |  | fusiform gyrus (19) | LH | -38 | -66 | -18 | 10 | 3,26 | 3,48 |
| Main effect of Education |  |  |  |  |  |  |  | Positive effect of Education H-L |  |  |  |  |  |  |  |
| claustrum | LH | -28 | 16 | -8 | 40 | 3,38 | 14,94 | claustrum | LH | -28 | 16 | -8 | 76 | 3,56 | 3,86 |
|  |  |  |  |  |  |  |  | supramarginal gyrus | RH | 60 | -56 | 22 | 28 | 3,48 | 3,76 |
|  |  |  |  |  |  |  |  | cingulate gyrus (31) | RH | 4 | -56 | 30 | 32 | 3,42 | 3,69 |
|  |  |  |  |  |  |  |  | middle frontal gyrus (9) | LH | -46 | 28 | 30 | 10 | 3,17 | 3,38 |
| Interaction Age x Education |  |  |  |  |  |  |  | Positive interaction YL/EH-YH/EL |  |  |  |  |  |  |  |
| superior frontal gyrus (8) | LH | -12 | 32 | 54 | 39 | 3,79 | 19,1 | superior frontal gyrus (8) | LH | -12 | 32 | 54 | 82 | 3,96 | 4,36 |
| parahippocampal gyrus | RH | 26 | -42 | 0 | 38 | 3,66 | 17,68 | parahippocampal gyrus | RH | 26 | -42 | 0 | 72 | 3,83 | 4,2 |
| claustrum | RH | 40 | -8 | 2 | 74 | 3,62 | 17,25 | claustrum | RH | 40 | -8 | 2 | 172 | 3,79 | 4,15 |
| insula (13) | RH | 42 | 2 | 8 | 74 | 3,32 | 14,43 | insula (13) | RH | 42 | 2 | 8 | 172 | 3,51 | 3,79 |
| superior temporal gyrus (39) | RH | 44 | -48 | 8 | 20 | 3,54 | 16,49 | superior temporal gyrus (39) | RH | 44 | -48 | 8 | 36 | 3,71 | 4,05 |
| hippocampus | LH | -30 | -46 | 4 | 10 | 3,38 | 14,89 | hippocampus | LH | -30 | -46 | 4 | 31 | 3,56 | 3,85 |
|  |  |  |  |  |  |  |  | middle frontal gyrus (6) | LH | 36 | -8 | 62 | 10 | 3,42 | 3,69 |
|  |  |  |  |  |  |  |  | insula (13) | LH | -34 | -4 | 16 | 17 | 3,29 | 3,53 |
|  |  |  |  |  |  |  |  | superior temporal gyrus (22) | LH | -50 | -16 | -6 | 15 | 3,26 | 3,49 |
|  |  |  |  |  |  |  |  | lentiform nucleus, putamen | RH | 30 | -4 | -12 | 11 | 3,23 | 3,46 |

<sup>a</sup> The education effect was found to involve the cerebellum using a more liberal threshold ( $p < 0.005$ uncorrected), in order to show an expected effect, considering the substantial impact of education on behavioral performance.

Regarding the percentage of signal change in both tasks, the elderly groups presented greater activation, and the younger ones greater deactivation. However, for phonemic fluency, the main effect of education meant that low-educated participants showed deactivation of the cerebellum, while the high-educated groups presented activation of this region. On the other hand, in the semantic task, the opposite pattern of activation was found: low-educated groups showed activation while high-educated groups presented deactivation of the left claustrum.

In light of correlations between the percentage of signal change and the total performance scores, some moderate to strong positive associations were found only for semantic VF. The percentage of signal change in the right claustrum tended to be correlated to semantic VF performance in HY adults (r = 0.527; p = 0.078). In addition, there was a correlation between the percentage of signal change in the right STG in LE participants (r = 0.682; p = 0.015). Finally, in this same region, a correlation was found in younger adults (r = 0.515; p = 0.01). Therefore, for these three associations, more deactivation may represent better performance.

## 4. Discussion

In this study, we aimed to describe the impact of age and education on behavioral performance in phonemic and semantic VF tasks, and the possibility of an interaction between these sociodemographic factors. We also sought to verify whether there were differences regarding neural activation and deactivation between age and education groups and whether this pattern correlated with their performance, by means of a mixed fMRI design. For the phonemic paradigm only, we found that education played a role: higher-educated adults performed better. On the other hand, for the semantic task, we observed an interaction between age and education: there was an education effect only for the younger group. In the fMRI findings, there were main effects of both age and education, as well as an interaction between them, in different brain areas depending on modality; sometimes the effect was seen in activation and in other cases it involved deactivation.

### 4.1 Behavioral findings

We found an interaction between age and education on semantic VF task performance, which is line with previous findings (36,39–42). In parallel, we found main effects of age and education on the phonemic VF task. Previous behavioral studies have reported similar findings about the effects of age (43–46) and education (22,47) on phonemic VF tasks. Differential effects of age and education depending on the type of VF paradigm are commonly reported in the literature (48,49) and may be related to the different brain circuits and cognitive functions (20,50) that are recruited more by each paradigm.

In addition, education effects were more frequent on behavioral assessment variables. This suggests that the sample of elderly people was a high-performing group. This finding is especially important regarding performance on the Vocabulary subtest (main effect of education only), which can be considered as an index of baseline semantic memory performance, from which VF can arise (51). Importantly, the Months task (control task) proved to be sensitive to the education factor, with LE adults performing worse than HE individuals. When analyzed together, these results indicate that the effects that were found can be attributed to the VF paradigm rather than to semantic memory or word generation activity alone.

### 4.2 fMRI findings per group

Given that, hitherto, no studies had presented the brain regions involved in phonemic and semantic VF tasks for four different age and education groups, the results of these first-level analyses may be very useful in developing a preliminary picture of the brain areas associated with the linguistic, mnestic and executive abilities underlying VF in HY, LY, HE and LE individuals. Although these brain regions have been investigated quite extensively in HY persons, and also quite frequently in HE adults, brain activation had not been sufficiently explored in either younger or older groups with less schooling. In light of the increasing recognition of the potential value of fMRI in clinical applications, studies that contribute evidence of activation patterns from normal participants should be helpful for the future development of normative scales evaluating individual patients’ outcomes (19).

In the phonemic VF paradigm, the right cingulate gyrus was among the most frequently activated regions for all groups except the HE group. This region has often been associated with VF performance (52). Its posterior portion has been related to attentional focus (53). The role of the insula, which was activated in the low-educated groups, has been increasingly explored, as it is associated with language production, repetition, complex linguistic processes and, more specifically, lexico-semantic associations (54). In view of the poorer performance associated with less education, and the role of the insula in language, this region may be related to language stimulation through formal educational experience. In line with this hypothesis, the claustrum was also important for the LY and LE groups; it is related to attentional control and segregation (55). For better-educated groups, two different brain regions proved important: the right cerebellum for HY people and the left middle frontal gyrus (MFG) for HE individuals. For the past three decades, the cerebellum has increasingly been found to be related to complex cognitive abilities (56), mainly language (57) and executive functions (58). Most VF fMRI studies have found that the left MFG plays an important role (see Costafreda et al. (20)).

In the semantic fluency paradigm, both LY and HY groups showed activation in the left caudate tail, whose role in working memory, mainly spatial (59), has already been recognized, probably due to the visual representations of each word through an image-based strategy. In accordance with the findings for phonemic VF, HE individuals also recruited the left MFG. This activation may be related to this region’s essential participation in switching between semantic subcategories (60). The cingulate gyrus also played a very important role for semantic VF in both elderly groups, as in Meinzer et al.’s (11) study, which found more pronounced activation in elderly participants only for a category fluency task.

A very interesting result refers to the right dominance of regions involved in both VF paradigms in the LE group. This is in line with the HAROLD model (12), which postulates greater right hemisphere activation in elderly people in order to try to maintain higher levels of performance, by recruiting right hemisphere brain regions to perform tasks normally accomplished by their left hemisphere homologues. One could also explain this result by means of Banich’s (61) theory that brain hemispheres couple and decouple to accomplish more cognitively demanding tasks. In this case, the task must have been more difficult for the LE population than for the other groups, therefore activating their right hemisphere more intensively to cope with a challenging task. This brain functional compensation strategy seemed to be necessary only for less-educated elderly participants, who seem to have been simultaneously impacted by their advanced age and lower schooling background. The concept of brain adaptation during aging has been widely explored, for instance through the scaffolding theory (62), which explains the contralateral activation as a function of inefficient inhibition related to behavioral deficits in elderly people. Cognitive deficits are more frequent in elderly participants who have less education. Thus, in an aging brain, compensation strategies may enhance recruitment of right hemisphere regions when age and education simultaneously play a role in cognition.

#### 4.2.1 Main effect of age on phonemic and semantic VF

Although age had no effect on performance in either task, it did affect activation in the left PCG and left STG in the phonemic VF task, as well as the left paracentral lobule and left PCG in the semantic paradigm. With regard to the left PCG, younger adults showed more deactivation in that area, while more activation was observed in elderly participants. Leech and Sharp (53) observed that this region deactivates more with attentionally demanding tasks, reinforcing the hypothesis that cognition is internally directed in younger people. In elderly subjects, the greater activation may be related to the left PCG’s role in memory retrieval (63,64). In addition, the left STG, which shows the same activation/deactivation pattern, has been associated with verbal working memory processing (65). For instance, even though they presented comparable performance, elderly scored lower scores in random number generation, which can be linked to compensation for working memory overload during the phonemic VF task for this group. Schneider-Garces et al. (66) revisited the CRUNCH model to predict activation differences between younger and elderly individuals and highlighted the hypothesis that greater brain activation can be observed in elderly people due to individual variations in working memory span. For a review, please refer to Park and Festini (67). Lastly, Meinzer et al. (11) found the paracentral lobule to be more activated in elderly subjects; this region has been associated with flexibility demands (68). Recently, Hoyau et al. (69) found that elderly use a neural compensation strategy for word retrieval involving the inferior frontal cortex and the medial temporal cortex.

#### 4.2.2 Main effect of education on phonemic and semantic VF

In line with the behavioral results, education also played a special role in brain activation and deactivation for both phonemic and semantic VF tasks. This finding is challenging to explain due to the lack of literature on the issue. In an fNIRS study, Heinzel et al. (70) noted that education played a greater role in phonemic VF than in a semantic task. In the phonemic paradigm in our study, we observed a main effect of education in the right cerebellum, with higher education associated with deactivation and the opposite pattern for less-educated subjects. Similarly, Bonnet et al. (71) used a Go/No-go fMRI paradigm and noted that healthy adults with higher education showed greater cerebellar activity than lower-educated people. Their main hypothesis was related to the cerebellum’s role in recruiting automatic strategies for more successful attentional – and we would add executive – processes. Furthermore, a group with right cerebellar lesions produced fewer words than a group with left cerebellar lesions and than healthy controls, showing this region’s important influence on phonemic word generation (72). Thus, it seems that the association between higher education and better performance in VF tasks, at least letter-based ones, may depend on the cerebellum and executive/attentional ability.

On the other hand, in the semantic VF modality, a main effect of years of education was found in the left claustrum. The pattern of activation and deactivation was the opposite of the one observed in the phonemic paradigm: in higher-educated subjects this region was deactivated, whereas it was activated in lower-educated adults. This pattern may be explained by compensation for low education by means of greater recruitment of attentional control during a continuous word search. In this regard, Smythies et al. (73) emphasized that the claustrum plays a special role during multicenter cognitive tasks, contributing to intracortical and intraclaustral synchronization. More specifically, then, one can predict that it may be needed for semantic association.

#### 4.2.3 Interaction between age and education on phonemic and semantic VF

Based on the results of behavioral studies, the interaction between age and education cannot be simply interpreted through our fMRI findings. Our preliminary hypothesis was that there may be a continuum of activation and deactivation corresponding to these four age and education groups. However, in the phonemic cue paradigm, we noticed a very interesting and surprising interaction in the left SOG: this area was deactivated more in the LY and HE groups than in the HY and LE groups. This region has been associated with a semantic network common to both words and images (74). This crossed effect of age and education in the under-recruitment of the left SOG may possibly be further explained in future studies if deactivation occurs in all groups to avoid a conflict of semantic strategies with a letter-based paradigm.

Conversely, in the semantic task an interaction between age and education was noted in the right claustrum, the STG and the bilateral hippocampus, representing greater activation in HE individuals than in the other groups; increased deactivation in these regions was observed in the following order in the other three groups: LE > HY > LY. The right claustrum, as mentioned in our discussion of first-level analyses, is closely related to attentional control. The hippocampus and parahippocampal gyrus have been linked to the process of categorical recovery (75). We hypothesized that high-educated elderly adults needed more expressive participation of attentional control.

Taken together, the results on activation and deactivation do not seem to present a simple linear relationship with performance in VF tasks. According to Binder (76), deactivation should be further analyzed in light of types of baseline tasks, especially in language tasks; thus, its nature has not yet been fully understood. Persson et al. (77) remarked that the deactivation response is less sensitive to demand in older subjects than in young adults. Reviewing the neuronal basis for task-negative responses, Linn et al. (78) explored two models: (1) greater activation (reductions in metabolic rate) associated with effortful tasks, and the opposite pattern (2) greater activation linked to less effort in cognitive activities. Thus, further studies should analyze the nature of the relationship between deactivation and performance in more depth in twofold linguistic and executive tasks.

### 4.3 Contributions and limitations

The novelty of this work was that it associated the less explored impact of education with the more frequently studied age effect on performance and brain neural correlates in a mixed phonemic and semantic VF design, with a 90-second paradigm and an automatic control task. The brain regions involved may be activated due to advanced age and/or lower educational background, and not necessarily due to clinical brain damage or brain dysfunction. In this way, our findings may be helpful in avoiding false fMRI positives, such as the widespread false positive findings of cognitive deficits.

Among caveats of the current study, we note that, although fluency in the months task has already been validated in the literature, such as by Birn et al. (7), considering the education effect, this task may have been too effortful for the less-educated participants. In addition, activation and deactivation patterns were quite weakly correlated with each group’s performance. Finally, further investigations should use larger samples. However, this was the first study to simultaneously consider two age groups and two educational groups, aiming to describe the brain regions involved in phonemic and semantic VF. More specifically, this study was the first to assess use of fMRI and an overt, long-duration, self-paced semantic and phonemic VF paradigm with lower-educated healthy adults.

### 4.4 Conclusions

The main aims of this study were twofold: to verify a potential impact of age and education or an interaction between these factors on phonemic and semantic fMRI VF; and (2) to examine possible neural correlates of these effects in brain area recruitment using a mixed-design fMRI VF task. Brain activation involved regions related to linguistic, executive and working memory abilities. Education was shown to a greater impact on performance than the age factor. Marsolais et al. (26) also posited that the age effect is less prominent than the education effect in VF, especially in highly educated samples. Moreover, the greater participation of the right hemisphere in coping with a high-demand task for older less-educated adults should be noted; it corroborates the literature on the role of cognitive reserve in aging, as well as the recruitment of additional brain areas to adapt to demanding tasks, as posited by Banich’s (61) model. Taken together, the behavioral and neuroimaging data provided by our study expand and advance existing knowledge of the interaction between age and education in a relevant linguistic task, correlating the behavioral results with the neural circuitry results.

Future studies should analyze brain and behavioral performance using another, even simpler, automatic baseline task for very low-educated populations, including illiterate participants. Another possible study could analyze in more depth the impact of stimulus complexity within both tasks (semantic and phonetic) on brain and behavioral performance. In addition, clinical populations could be investigated in comparison to healthy controls. Considering that VF is one of the most sensitive and effective tools for clinical neurocognitive diagnosis, further studies should shed more light on our understanding of all the sociodemographic and cognitive factors that impact VF ability.

## References

1. Wagner S, Sebastian A, Lieb K, Tüscher O, Tadić A. A coordinate-based ALE functional MRI meta-analysis of brain activation during verbal fluency tasks in healthy control subjects. BMC Neurosci. 2014;15:19.

2. Joanette Y, Ska B, Côté H. Protocole Montréal d’Evaluación de la Communication. Montréal: Ortho Editions; 2004.

3. Zimmermann N, Branco L, Ska B, Gasparetto EL, Joanette Y, Fonseca R. Verbal Fluency in Right Brain Damage: Dissociations Among Production Criteria and Duration. Appl Neuropsychol Adult. 2013;(March 2014):1–9.

4. Heim S, Eickhoff SB, Amunts K. Specialisation in Broca’s region for semantic, phonological, and syntactic fluency? Neuroimage. 2008;40(3):1362–8.

5. Shao Z, Janse E, Visser K, Meyer AS. What do verbal fluency tasks measure? Predictors of verbal fluency performance in older adults. Front Psychol. 2014 Jul;5(JUL):1–10.

6. Zimmermann N, Cardoso CDO, Trentini CM, Grassi-Oliveira R, Fonseca RP. Brazilian preliminary norms and investigation of age and education effects on the Modified Wisconsin Card Sorting Test, Stroop Color and Word test and Digit Span test in adults. Dement Neuropsychol. 2015;9(2):120–7.

7. Birn RM, Kenworthy L, Case L, Caravella R, Jones TB, Bandettini PA, et al. NeuroImage Neural systems supporting lexical search guided by letter and semantic category cues : A self-paced overt response fMRI study of verbal fluency. Neuroimage. 2010;49(1):1099–107.

8. Baciu M, Boudiaf N, Cousin E, Perrone-Bertolotti M, Pichat C, Fournet N, et al. Functional MRI evidence for the decline of word retrieval and generation during normal aging. Age (Dordr) [Internet]. 2015/12/28. 2016 Feb;38(1):3. Available from: https://pubmed.ncbi.nlm.nih.gov/26711670

9. Marsolais Y, Perlbarg V, Benali H, Joanette Y. Age-related changes in functional network connectivity associated with high levels of verbal fluency performance. Cortex. 2014;58:123–38.

10. Meinzer M, Wilser L, Flaisch T, Eulitz C, Rockstroh B, Conway T, et al. NIH Public Access. 2009;21(10):2007–18.

11. Meinzer M, Wilser L, Flaisch T, Eulitz C, Rockstroh B, Conway T, et al. Neural signatures of semantic and phonemic fluency in young and old adults. J Cogn Neurosci. 2009;21(10):2007–18.

12. Cabeza R. Hemispheric Asymmetry Reduction in Older Adults : The HAROLD Model. Psychol Aging. 2002;17(1):85–100.

13. Reuter-Lorenz PA, Campbell KA. Neurocognitive ageing and the Compensation Hypothesis. Curr Dir Psychol Sci. 2008;17(3):177–82.

14. Dehaene S, Cohen L, Morais J, Kolinsky R. Illiterate to literate: behavioural and cerebral changes induced by reading acquisition. Nat Rev Neurosci. 2015;16(4):234–44.

15. Stern Y. Cognitive reserve. Neuropsychologia. 2009;47(2009):2015–28.

16. Beausoleil N, Fortin R, Le Blanc B, Joanette Y. Unconstrained oral naming performance in right- and left-hemisphere-damaged individuals: When education overrides the lesion. Aphasiology. 2003;17(2):143–58.

17. Nagels A, Kircher T, Dietsche B, Backes H, Marquetand J, Krug A. Neural processing of overt word generation in healthy individuals: the effect of age and word knowledge. Neuroimage. 2012 Jul;61(4):832–40.

18. Kim J, Chey J, Kim SE, Kim H. The effect of education on regional brain metabolism and its functional connectivity in an aged population utilizing positron emission tomography. Neurosci Res. 2015;94:50–61.

19. Allen MD, Fong AK. Clinical application of standardized cognitive assessment using fMRI. II. verbal fluency. Behav Neurol. 2008;20(3–4):141–52.

20. Costafreda SG, Fu CHY, Lee L, Everitt B, Brammer MJ, David AS. A systematic review and quantitative appraisal of fMRI studies of verbal fluency: role of the left inferior frontal gyrus Hum Brain Mapp. 2006 Oct;27(10):799–810.

21. Abrahams S, Goldstein LH, Simmons A, Brammer MJ, Williams SCR, Giampietro VP, et al. Functional magnetic resonance imaging of verbal fluency and confrontation naming using compressed image acquisition to permit overt responses. Hum Brain Mapp. 2003 Sep;20(1):29–40.

22. Aziz KA, Khater MS, Emara T, Tawfik HM, Rasheedy D, Mohammedin AS, et al. Effects of age, education, and gender on verbal fluency in healthy adult Arabic-speakers in Egypt. Appl Neuropsychol Adult. 2016;9095(June):1–11.

23. de Azeredo Passos VM, Giatti L, Bensenor I, Tiemeier H, Ikram MA, de Figueiredo RC, et al. Education plays a greater role than age in cognitive test performance among participants of the Brazilian Longitudinal Study of Adult Health (ELSA-Brasil). BMC Neurol. 2015;15(1):191.

24. Nielsen TR, Waldemar G. Effects of literacy on semantic verbal fluency in an immigrant population. Neuropsychol Dev Cogn B Aging Neuropsychol Cogn. 2016;5585(March):1–13.

25. van der Elst W, van Boxtel MPJ, van Breukelen GJP, Jolles J. Normative data for the Animal, Profession and Letter M Naming verbal fluency tests for Dutch speaking participants and the effects of age, education, and sex. J Int Neuropsychol Soc. 2006;12(1):80–9.

26. Marsolais Y, Methqal I, Joanette Y. Marginal neurofunctional changes in high-performing older adults in a verbal fluency task. Brain Lang. 2015;140:13–23.

27. Brito GN, Brito LS, Paumgartten FJ, Lins MF. Lateral preferences in Brazilian adults: an analysis with the Edinburgh Inventory. Cortex. 1989;25:403–15.

28. Folstein MF, Folstein SE, McHugh PR. “Mini-mental state”: A practical method for grading the cognitive state of patients for the clinician. J Psychiatr Res. 1975;12(3):189–98.

29. Kochhann R, Varela JS, Lisboa CSDM, Chaves MLF. The Mini Mental State Examination Review of cutoff points adjusted for schooling in a large Southern Brazilian sample. Dement Neuropsychol. 2010;4(1):35–41.

30. Lopez MN, Charter R a, Mostafavi B, Nibut LP, Smith WE. Psychometric properties of the Folstein Mini-Mental State Examination. Assessment. 2005 Jun;12(2):137–44.

31. Beck AT, Steer RA, Brown GK. BDI-II Inventário de Depressão de Beck. 1st ed. São Paulo: Pearson/Casa do Psicólogo; 2012.

32. Gorenstein C, Wang YP, Argimon IL, Werlang BSG. Manual do Inventário de Depressão de Beck - BDI-II Adaptação Brasileira. 1^a^ Edição. São Paulo: Casa do Psicólogo; 2011. 156 p.

33. Wechsler D. Escala Wechsler de Inteligência para Adultos (WAIS-III). São Paulo: Pearson/Casa do Psicólogo; 2004.

34. Nascimento E. WAIS-III: Escala de Inteligência Wechsler para Adultos - manual técnico. Wechsler D, editor. São Paulo: Casa do Psicólogo; 2004.

35. Fonseca RP, Parente alice de MP, Cote H, Ska B, Joanette Y. Bateria MAC - Bateria Montreal de Avaliação da Comunicação (instrumento completo de avaliação). Pró-Fono. Andrade CRF de, Mansur LL, Vilanova LCP, editors. 2008.

36. Gaillard WD, Sachs BC, Whitnah JR, Ahmad Z, Balsamo LM, Petrella JR, et al. Developmental aspects of language processing: fMRI of verbal fluency in children and adults. Hum Brain Mapp. 2003 Mar;18(3):176–85.

37. Worsley KJ, Liao CH, Aston J, Petre V, Duncan GH, Morales F, et al. A general statistical analysis for fMRI data. Neuroimage. 2002;15:1–15.

38. Sun X, Briel M, Walter SD, Guyatt GH. Is a subgroup effect believable? Updating criteria to evaluate the credibility of subgroup analyses. Br Med J. 2010;340(mar30 3):c117–c117.

39. Brucki SMD, Rocha MSG. Category fluency test: Effects of age, gender and education on total scores, clustering and switching in Brazilian Portuguese-speaking subjects. Brazilian J Med Biol Res. 2004;37(12):1771–7.

40. Ostrosky-Solis F, Gutierrez AL, Flores MR, Ardila A. Same or different? Semantic verbal fluency across Spanish-speakers from different countries. Arch Clin Neuropsychol. 2007;22(3):367–77.

41. Stricks L, Pittman J, Jacobs DM, Sano M, Stern Y. Normative data for a brief neuropsychological battery administered to English- and Spanish-speaking community-dwelling elders. J Int Neuropsychol Soc. 1998;4:311–318.

42. Taporoski TP, Duarte NE, Pompéia S, Sterr A, Gómez LM, Alvim RO, et al. Heritability of semantic verbal fluency task using time-interval analysis. PLoS One [Internet]. 2019 Jun 11;14(6):e0217814–e0217814. Available from: https://pubmed.ncbi.nlm.nih.gov/31185027

43. Bryan J, Luszcz M a, Crawford JR. Verbal knowledge and speed of information processing as mediators of age differences in verbal fluency performance among older adults. Psychol Aging. 1997;12(3):473–8.

44. Cavaco S, Gonçalves A, Pinto C, Almeida E, Gomes F, Moreira I, et al. Semantic fluency and phonemic fluency: regression-based norms for the Portuguese population. Arch Clin Neuropsychol. 2013 May;28(3):262–71.

45. Lanting S, Haugrud N, Crossley M. The effect of age and sex on clustering and switching during speeded verbal fluency tasks. J Int Neuropsychol Soc. 2009 Mar;15(2):196–204.

46. Hatta, Hotta, Kato, Fujiwara, Iwahara. Dissociation in Age-Related Developmental Trajectories Between Phonetic Fluency and Semantic Fluency Tests: Analysis of Longitudinal Data From the Yakumo Study. Am J Psychol. 2020 Jul 1;133:197.

47. Buriel Y, Gramunt-Fombuena N, Böhm P, Rodés E, Peña-Casanova J. Fluencia verbal: Estudio normativo piloto en una muestra española de adultos jóvenes (20 a 49 años). Neurología. 2004;19(4):153–9.

48. Mathuranath PS, George A, Cherian PJ, Alexander A, Sarma SG, Sarma PS. Effects of age, education and gender on verbal fluency. J Clin Exp Neuropsychol. 2003;25(8):1057–64.

49. Zimmermann N, Alice M, Pimenta DM, Joanette Y, Paz R. Unconstrained, Phonemic and Semantic Verbal Fluency : Age and Education Effects, Norms and Discrepancies. Psicol Reflex e Crit. 2014;27(1):1–9.

50. Brickman AM, Paul RH, Cohen RA, Williams LM, MacGregor KL, Jefferson AL, et al. Category and letter verbal fluency across the adult lifespan: Relationship to EEG theta power. Arch Clin Neuropsychol. 2005;20(5):561–73.

51. Ruff RM, Light RH, Parker SB, Levin HS. The psychological construct of word fluency. Brain Lang. 1997;(57):394–405.

52. Halari R, Sharma T, Hines M, Andrew C, Simmons A, Kumari V. Comparable fMRI activity with differential behavioural performance on mental rotation and overt verbal fluency tasks in healthy men and women. Exp Brain Res. 2006;169(1):1–14.

53. Leech R, Sharp DJ. The role of the posterior cingulate cortex in cognition and disease. Brain. 2014;137(1):12–32.

54. Ardila A, Bernal B, Rosselli M. Participation of the insula in language revisited: A meta-analytic connectivity study. J Neurolinguistics. 2014;29(November 2015):31–41.

55. Goll Y, Atlan G, Citri A. Attention: the claustrum. Trends Neurosci. 2015;38(8):486–95.

56. Grossauer S, Koeck K, Kau T, Weber J, Vince GH. Behavioral disorders and cognitive impairment associated with cerebellar lesions. J Mol psychiatry. 2015;3(1):5.

57. Dobrianskyj Weber LN, Leite CR, Stasiak GR, Da Silva Santos CA, Forteski R. O estresse no trabalho do professor. Imagens da Educ [Internet]. 2015 Nov 12 [cited 2020 Aug 15];5(3):40. Available from: http://dx.doi.org/10.4025/imagenseduc.v5i3.25789

58. Nowrangi M a, Lyketsos C, Rao V, Munro C a. Systematic review of neuroimaging correlates of executive functioning: converging evidence from different clinical populations. J Neuropsychiatry Clin Neurosci. 2014;26:114–25.

59. Postle BR, D’Esposito M. Dissociation of human caudate nucleus activity in spatial and nonspatial working memory: an event-related fMRI study. Cogn Brain Res. 1999 Jul;8(2):107–15.

60. Hirshorn EA, Thompson-Schill SL. Role of the left inferior frontal gyrus in covert word retrieval: Neural correlates of switching during verbal fluency. Neuropsychologia. 2006;44(12):2547–57.

61. Banich MT. Interaction between the hemispheres and its implications for the processing capacity of the brain. In: The asymmetrical brain. Cambridge, MA, US: MIT Press; 2003. p. 261–302.

62. Cramer SC, Sur M, Dobkin BH, O’Brien C, Sanger TD, Trojanowski JQ, et al. Harnessing neuroplasticity for clinical applications. Brain. 2011;134(6):1591–609.

63. Maddock RJ, Garrett AS, Buonocore MH. Remembering familiar people: The posterior cingulate cortex and autobiographical memory retrieval. Neuroscience. 2001;104(3):667–76.

64. Qin S, Piekema C, Petersson KM, Han B, Luo J, Fernández G. Probing the transformation of discontinuous associations into episodic memory: An event-related fMRI study. Neuroimage. 2007;38(1):212–22.

65. Buchsbaum B, Hickok G, Humphries C. Role of left posterior superior temporal gyrus in phonological processing for speech perception and production. Cogn Sci. 2001;25(5):663–78.

66. Schneider-Garces NJ, Gordon BA, Brumback-Peltz CR, Shin E, Lee Y, Sutton BP, et al. Span, CRUNCH, and Beyond: Working Memory Capacity and the Aging Brain. J Cogn Neurosci. 2010 Apr;22(4):655–69.

67. Park DC, Festini SB. Theories of Memory and Aging: A Look at the Past and a Glimpse of the Future. Journals Gerontol Ser B Psychol Sci Soc Sci. 2016;00(00):gbw066.

68. Niendam TA, Laird AR, Ray KL, Dean YM, Carter CS. NIH Public Access. 2013;12(2):241–68.

69. Hoyau E, Roux-Sibilon A, Boudiaf N, Pichat C, Cousin E, Krainik A, et al. Aging modulates fronto-temporal cortical interactions during lexical production. A dynamic causal modeling study. Brain Lang [Internet]. 2018;184:11–9. Available from: http://www.sciencedirect.com/science/article/pii/S0093934X17300937

70. Heinzel S, Metzger FG, Ehlis A-C, Korell R, Alboji A, Haeussinger FB, et al. Aging-related cortical reorganization of verbal fluency processing: a functional near-infrared spectroscopy study. Neurobiol Aging. 2012;34(2):439–50.

71. Bonnet MC, Dilharreguy B, Allard M, Deloire MSA, Petry KG, Brochet B. Differential cerebellar and cortical involvement according to various attentional load: Role of educational level. Hum Brain Mapp. 2009;30(4):1133–43.

72. Schweizer TA, Alexander MP, Susan Gillingham BA, Cusimano M, Stuss DT. Lateralized cerebellar contributions to word generation: A phonemic and semantic fluency study. Behav Neurol. 2010;23(1–2):31–7.

73. Smythies JR, Edelstein LR, Ramachandran VS. Hypotheses Relating to the Function of the Claustrum. Claustrum Struct Funct Clin Neurosci. 2014;6(August):299–352.

74. Vandenberghe R, Price C, Wise R, Josephs O, Frackowiak RS. Functional anatomy of a common semantic system for words and pictures. Vol. 383, Nature. 1996. p. 254–6.

75. Pihlajamäki M, Tanila H, Hänninen T, Könönen M, Laakso M, Partanen K, et al. Verbal fluency activates the left medial temporal lobe: a functional magnetic resonance imaging study. Ann Neurol. 2000;47(4):470–6.

76. Binder JR. Task-induced deactivation and the “resting” state. Neuroimage. 2012;62(2):1086–91.

77. Persson J, Lustig C, Nelson JK, Reuter-Lorenz PA. Age differences in deactivation: a link to cognitive control? J Cogn Neurosci. 2007 Jun;19(6):1021–32.

78. Linn MC. Designing Standards for Lifelong Science Learning. J Eng Educ [Internet]. 2010 Apr 1;99(2):103–5. Available from: https://doi.org/10.1002/j.2168-9830.2010.tb01047.x

